# Spligation enables programmable chimeric RNA generation in living cells

**DOI:** 10.64898/2026.03.06.709984

**Authors:** David A. Colognori, Kevin M. Wasko, Marena I. Trinidad, Zehan Zhou, Jennifer A. Doudna

## Abstract

The ability to precisely modify RNA offers opportunities to manipulate the flow of genetic information and influence transcript stability, localization and translation. RNA-targeting technologies enable RNA knockdown, base editing and *trans*-splicing, but more extensive transcript changes typically require genome editing or rely on the endogenous splicing machinery. Based on the ability of type III-A CRISPR-Csm complexes to catalyze programmable RNA cleavage in human cells, we investigated their potential to induce site-specific deletions while leaving the remainder of the transcript intact. Our data show that CRISPR-Csm complexes can generate short and long RNA excisions within a target transcript, and that the efficiency of this process is enhanced by fusion of Csm to the RNA ligase RtcB. Furthermore, cleavage of two different transcripts can trigger subsequent *trans*-ligation of the cleaved products into a chimeric transcript (“spligation”). Finally, we apply spligation to endogenous transcripts, using Csm to generate recombinant mRNA in cells independent of canonical splice sites. Collectively, this approach enables new forms of precise RNA manipulation in cells with potential applications in human disease.

## Introduction

CRISPR-based technologies have revolutionized genome engineering by enabling programmable nucleic acid manipulation^1^. RNA-guided enzymes including Cas9 and Cas12 allow precise cleavage of double-stranded DNA, which can be leveraged to introduce targeted genetic changes^2^. Such tools have opened new avenues for therapeutic genome editing and deepened our understanding of DNA repair mechanisms. These advances and the identification of RNA-targeting CRISPR systems have raised the possibility of using analogous approaches to induce targeted changes in cellular transcripts for both fundamental research and therapeutic applications.

Targeted RNA cleavage has been demonstrated using type VI RNA-targeting CRISPR-Cas13 nucleases, enabling destruction of cellular RNAs^3^. In addition, however, Cas13-mediated RNA cleavage also triggers widespread nonspecific cellular RNA degradation^4,5^. As a result, Cas13 has been incapable of introducing defined deletions in target RNAs with the precision and versatility achieved with Cas9-mediated DNA editing. Catalytically inactive versions of Cas13 can induce exon skipping and *trans*-RNA splicing^6^, but these methods are restricted to naturally occurring splice sites. The type III-E Cas7-11 nuclease, which cuts only once at the target site without *trans*-cleavage activity, exhibits low cleavage efficiency and poor specificity in human cells^7^. Expanding the RNA editing toolkit thus holds value to enable precise transcript manipulation for both research and possible therapeutic use.

CRISPR-Csm, an RNA-guided RNA-targeting protein complex, can induce site-specific deletions of cellular transcripts. Csm generates RNA cleavage products bearing 2′,3′-cyclic phosphate and 5′-hydroxyl termini that remain bound to Csm for hours after the release of internal RNA fragments^8^. RtcB is an evolutionarily conserved RNA ligase involved in tRNA maturation^9^ and unconventional XBP1 splicing^10^ that specifically recognizes and joins transcripts with the same end chemistries generated by Csm. Previously, RtcB was shown to ligate RNA fragments produced by Csm cleavage *in vitro*^10^, and was recently found to be responsible for RNA ligation following Csm-mediated cleavage in human cells^7^.

We show here that CRISPR-Csm-mediated cleavage of two RNA molecules coupled with RtcB ligation can generate chimeric transcripts in a process we term spligation. Co-localization of these enzymatic activities with a Csm-RtcB fusion construct enhances spligation efficiency. Alternative methods of generating 5′-hydroxyl termini in donor transcripts further enhanced the efficiency of spligation. This optimized technique was used to introduce kilobase-scale additions to endogenous mRNA transcripts in human cells unrestricted by canonical splice sites. Together, these results demon-strate the utility of targeted transcript alterations to test and control cellular RNA functions with spligation.

## Results

### CRISPR-Csm-mediated RNA breaks are re-ligated in human cells

We recently harnessed the type III-A CRISPR-Csm complex from *Streptococcus thermophilus* as a tool for programmable RNA cleavage in human cells^11^ (Fig. 1A). These experiments showed that nuclear-localized Csm complexes cleave transcripts complementary to a Csm-associated crRNA, leading to degradation of >90% of target transcripts (Fig. 1B, C). When examining the sequences of the remaining transcripts, we noticed a small fraction that appeared to have been cleaved and religated rather than degraded, in agreement with recent observations made using an identical CRISPR-Csm system in HEK293T cells^7^. Re-ligated transcripts contain a characteristic pattern of missing nucleotides in increments of 6, corresponding to the nuclease cut sites in the assembled Csm complex (Fig. 1D. Supplementary Table 1). This pattern was observed for two unique crRNAs targeting the noncoding RNA XIST (XIST crRNA 2 shown as example; XIST crRNA 1 in Supplementary Table 2). 12% of residual XIST RNA contained short deletions spanning precisely 6, 12, 18 or 24 nucleotides (nts). The major species contained deletions of 6 nucleotides within the innermost cleavage segments, consistent with prior biochemical characterization of Csm activity *in vitro*^12^.

**Figure 1:**
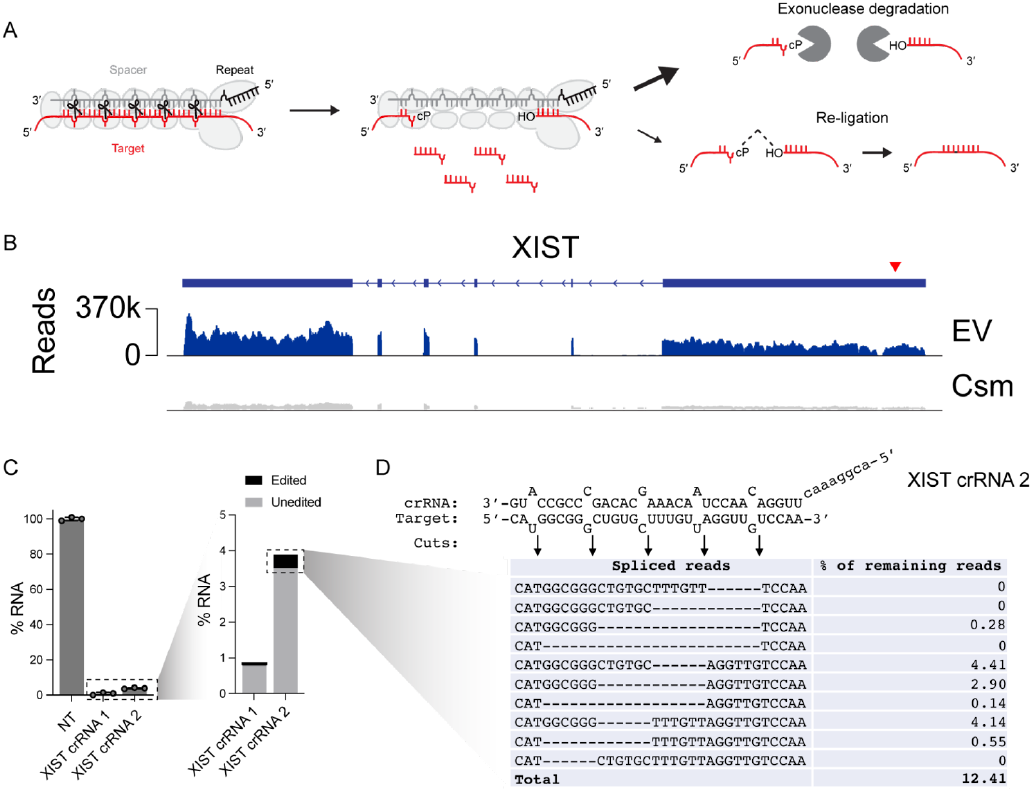
CRISPR-Csm-induced RNA breaks can be re-ligated in human cells. **(A)** Cartoon of Csm complex bound to crRNA and target RNA. Cleavage occurs in increments of 6 nt along the target, leaving behind a 2′,3′-cylclic phosphate (cP) and 5′-hydroxyl. **(B)** RNA-seq data showing Csm-mediated cleavage mostly leads to transcript-wide depletion compared to empty vector (EV) control. Cleavage site indicated by red arrowhead. **(C)** RT-PCR showing Csm-mediated cleavage of XIST RNA using two different crRNAs leads to >95% RNA depletion in HEK293T cells. Error bars indicate mean ± s.d. of three biological replicates. **(D)** Following Csm-mediated cleavage, ′12% of remaining transcripts contain small deletions that align with Csm’s cleavage pattern.

### Csm-RtcB fusions improve efficiency of RNA ligation and premature stop codon removal

To improve ligation efficiency following RNA cleavage in cells, we first validated a reporter system (CMV-mCherry-stop-eGFP) that would result in eGFP fluorescence following excision of a premature stop codon (Fig. 2A). We targeted the Csm complex to the stop codon, testing several individual crRNAs that recognize sequences tiled across the stop codon region (Fig. S1A). Because we expect most cut transcripts to be degraded, the tandem reporter system allows monitoring of both RNA cleavage (mCherry knockdown) and ligation (eGFP gain) simultaneously. Indeed, following transfection of HEK293T cells, only complexes programmed with a stop codon-targeting crRNA led to gain of eGFP fluorescence, as observed by fluorescence microscopy and flow cytometry (Fig. 2B, C). To assay editing directly at the RNA level, we extracted RNA from Csm-transfected cells and performed RT-PCR across the edited reporter locus, sequencing of which verified stop codon removal (Fig. 2A, D, Supplementary Table 3).

**Figure 2:**
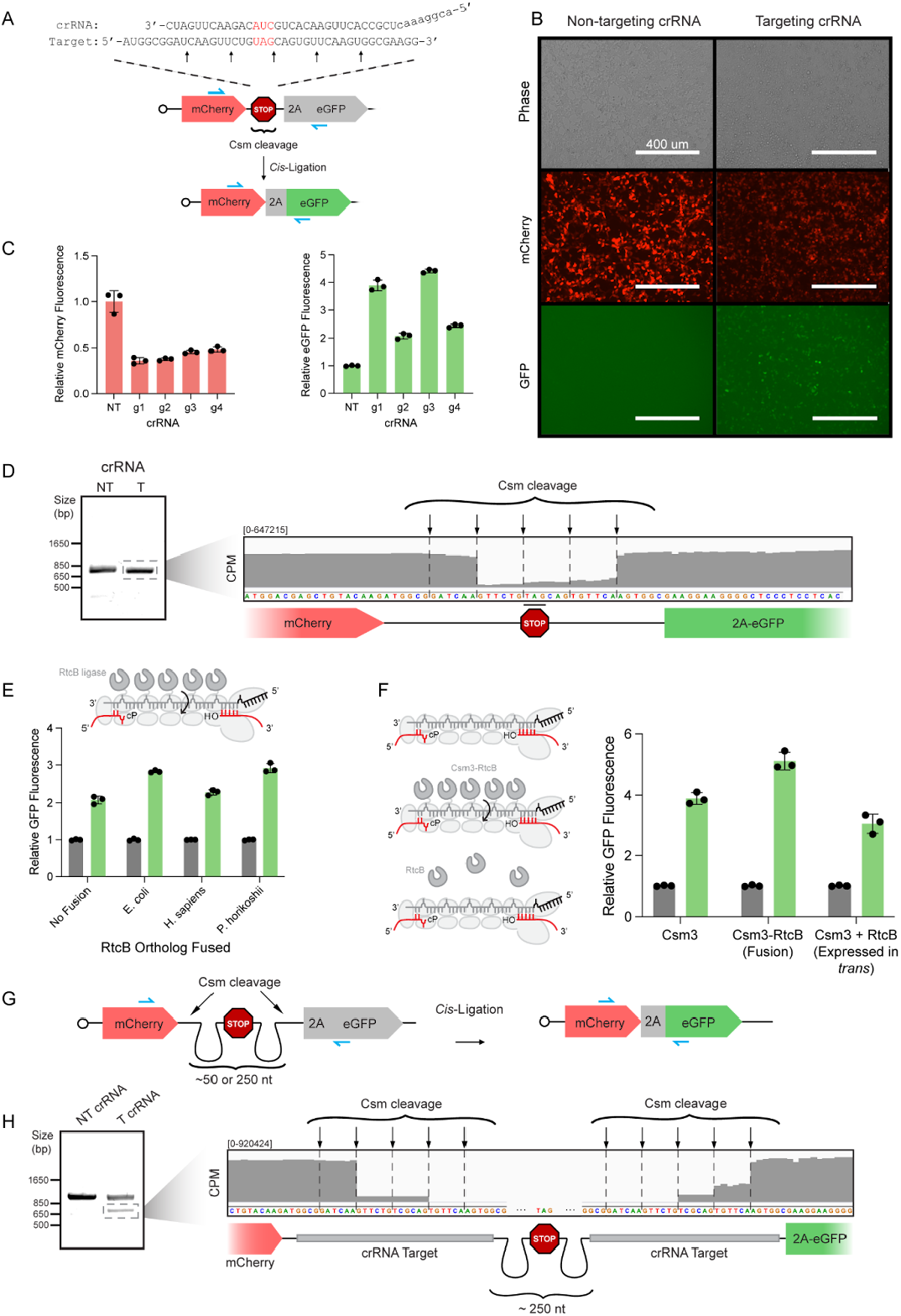
Csm-RtcB fusions result in increased re-ligation following transcript cleavage. **(A)** Schematic of CMV-mCherry-stop-eGFP reporter construct. Csm cleavage around the stop codon and subsequent re-ligation leads to stop codon removal while maintaining downstream reading frame, causing expression of eGFP. Zoom-in of Csm target site shows the positioning of an example crRNA. Stop codon is highlighted in red. Black and grey arrows show cleavage sites for crRNAs 1/3 and crRNAs 2/4, respectively. Blue arrows show location of PCR primers used in (D). **(B)** Brightfield and fluorescent microscopy images of HEK293T cells introduced with reporter, Csm, and either non-targeting or targeting crRNA. Red channel shows decreased mCherry signal following Csm cleavage while green channel shows appearance of eGFP signal following RtcB re-ligation. Scale bar is 400 um. **(C)** Flow cytometry quantification of mean fluorescence intensities of mCherry (left) and GFP (right) in transfected cells across tiled crRNAs normalized to the non-targeting crRNA control condition. **(D)** RNA was extracted from cells in (B,C), reverse-transcribed into cDNA, and PCR amplified using the primers indicated in blue in (A). The PCR product was resolved on a 1.5% agarose gel and the indicated band was excised and sequenced. Inset shows sequencing coverage, in counts per million (CPM), across the Csm cleavage site. Reporter sequence and diagram depicted below. Strategy of fusing RtcB ligase to Csm3 subunits to improve ligation efficiency through proximity. Flow cytometry quantification of GFP mean fluorescence intensities in transfected cells across Csm3-RtcB ortholog fusions normalized to the unfused Csm3 control. **(F)** Schematic of the Csm complex alone, with Csm3-RtcB fusions, or with RtcB expressed from a separate cassette (left). Flow cytometry quantification of GFP mean fluorescence intensities in transfected cells across conditions normalized to the unfused Csm3 control. **(G)** Schematic of CMV-mCherrycut-stop-cut-eGFP (dual cleavage) reporter construct. Csm cleavage on both sides of stop codon and subsequent re-ligation in cis is needed for downstream of expression of eGFP. Blue arrows show the location of PCR primers used in (H). **(H)** RNA was extracted from cells transfected with the construct shown in (G) with Csm target sites spaced ′250nt apart, reverse-transcribed into cDNA, and PCR amplified using the primers indicated in blue in (G). The PCR product was resolved on a 1.5% agarose gel and the indicated band excised and sequenced. Inset shows sequencing coverage across the Csm cleavage sites. Reporter sequence and diagram depicted below. Error bars indicate mean ± s.d. of three biological replicates.

We next created fusion constructs that physically link RtcB to the Csm complex. Based on a prior demonstration that fusing eGFP to Csm3 (the most abundant Csm complex subunit) enables live-cell RNA imaging^13^, a similar strategy was used to fuse human, archaeal (*Pyrococcus horikoshii*) or bacterial (*Escherichia coli*) RtcB to the C-terminus of Csm3 (Fig. 2E).

Interestingly, in our reporter system, human RtcB had no effect on eGFP fluorescence, but *E. coli* RtcB led to a ′30% increase compared to Csm alone (Fig. 2E). We screened a variety of linkers to fuse RtcB to either the N- or C-terminus of Csm3 to optimize for re-ligation efficiency (Fig. S1B). Over-expression of RtcB in *trans* was insufficient to improve ligation efficiency, suggesting that colocalization of the endonuclease and ligase is key (Fig. 2F). The increased performance of bacterial RtcB compared to its human counterpart could be due to its multi-turnover enzymatic activity, unlike human RtcB which is a single-turnover enzyme in the absence of its cofactor Archease^9^. We noticed that while the archeal RtcB fusion increased eGFP fluorescence, it only weakly reduced mCherry expression, similar to the human RtcB fusion construct (Fig. S1C). We speculate that the larger size of these orthologs may interfere with Csm complex assembly or function.

Having observed stop-codon removal, we next wondered whether targeting two sites in a transcript with CRISPR-Csm would enable excision of longer stretches of RNA. To test this, we constructed a second reporter gene containing a stop codon flanked by two identical Csm cleavage sites ′50 nts apart (CMV-mCherry-cut-stop-cut-eGFP) (Fig. 2G). Only cleavage at both sites and subsequent removal of the intervening stop region enables eGFP expression. RT-PCR analysis of the edited region of the reporter transcript revealed a truncated product, which we verified to be the reporter with the stop codon region removed (Fig. S1D, Supplementary Table 4). We repeated these experiments using a similar reporter construct with the Csm cleavage sites placed even further (′300 nts) apart and observed similar results as above (Fig. 2G, H, Supplementary Table 5). These results demonstrate that RtcB fusions enhance religation following Csm cleavage and that Csm is capable of inducing targeted RNA excisions across a wide range of excision lengths.

### Csm enables programmable ligation of RNA molecules in *trans*

Though our two-cut reporter was designed to monitor large deletions within a given transcript, it cannot be determined whether the resulting edits arise from *cis*-ligation or *trans*-ligation between two otherwise identical transcripts. Therefore, we next tested whether Csm-mediated cleavage of two separate transcripts could lead to ligation in *trans. Trans*-ligation of RNA mediated by endogenous RtcB was recently reported using self-cleaving ribozymes, which produce end chemistries identical to those generated by Csm cleavage^14^. However, transcripts compatible with the ribozyme-based system must be provided exogenously, whereas Csm can be programmed to target endogenous RNAs. To test the possibility of Csm-driven *trans*-ligation, we split eGFP into N- and C-terminal halves according to previous methods^15^, each encoded by separate plasmids. If Csm-mediated cleavage of both transcripts enables *trans*-ligation, it would lead to reconstitution of eGFP (Fig. 3A).

**Figure 3:**
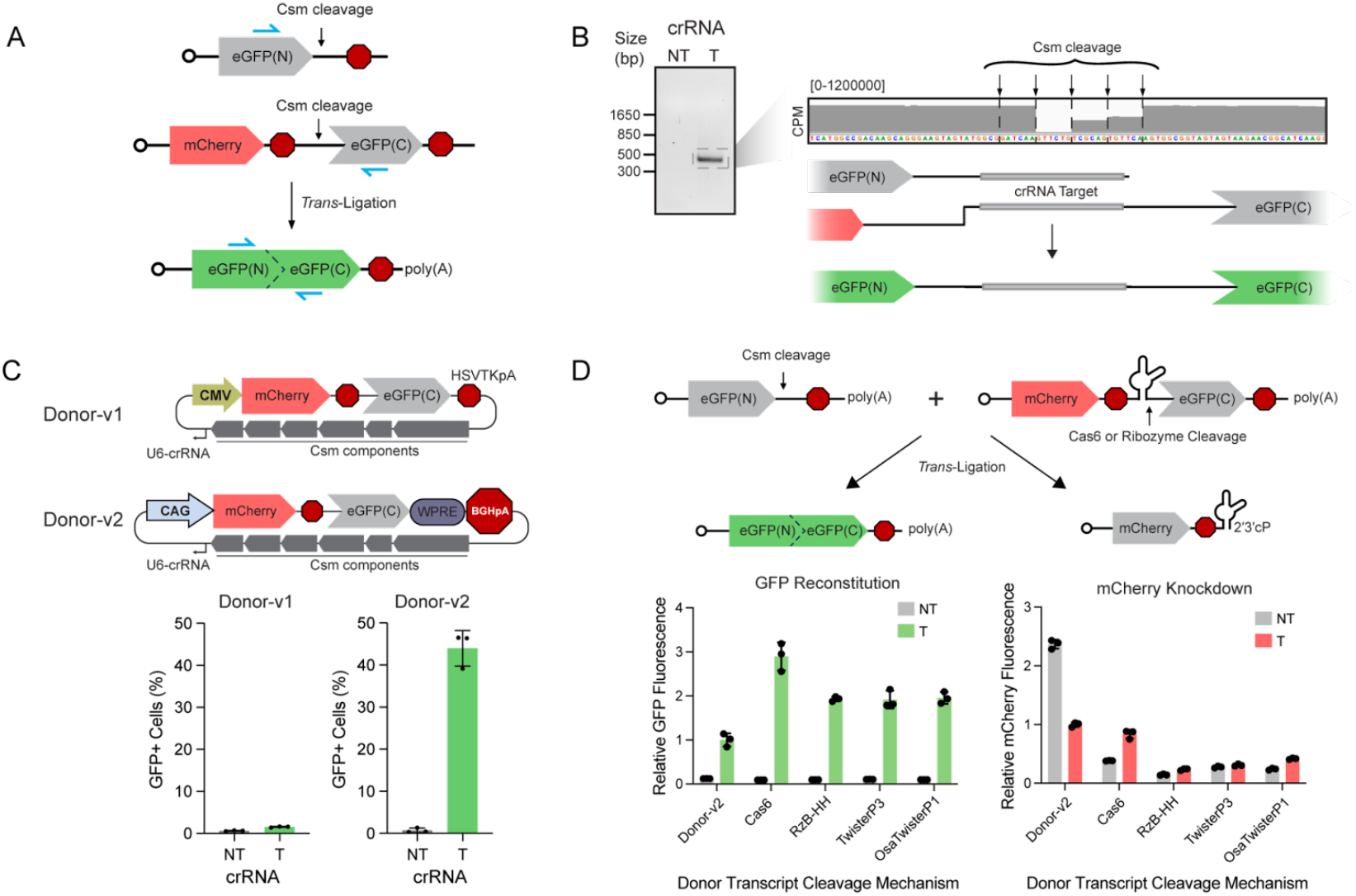
Csm-targeted trancripts can be ligated in *trans* to donor transcripts bearing a 5’-hydroxyl terminus. **(A)** Cartoon of CMV-eGFP(N) / CMV-mCherry-eGFP(C) (bipartite) reporter construct. Csm cleavage of both transcripts and subsequent re-ligation in trans is needed to restore expression of full-length eGFP. Blue arrows show the location of PCR primers used in (B). **(B)** RNA was extracted from cells transfected with the constructs outlined in (A), reverse-transcribed into cDNA, and PCR amplified using the primers indicated in blue in (A). The PCR product was resolved on a 1.5% agarose gel and the indicated band excised and sequenced. Inset shows sequencing coverage across the Csm cleavage sites. Reporter sequences and diagram depicted below. **(C)** Schematics of v1 and v2 trans-ligation reporter plasmids (top) and resulting GFP fluorescence observed in transfected cells as a percentage of the total mCherry+ population (bottom). **(D)** Schematic of v3 trans-ligation system using alternative 5′-hydroxyl generation mechanisms for the donor transcript (top). Flow cytometry quantification of mean fluorescence intensities of GFP (left) and mCherry (right) in transfected cells across tested conditions normalized to the Donor-v2 condition (bottom). Error bars indicate mean ± s.d. of three biological replicates.

We observed a rare population of cells (′1%) that exhibited eGFP fluorescence following transfection with plasmids encoding Csm-RtcB, the bipartite reporter (Donor-v1), and a targeting crRNA, versus the non-targeting crRNA condition (Fig S2B). To confirm *trans*-ligation, we extracted RNA and performed RT-PCR across the predicted junction. A product of the expected size was observed only for the targeting crRNA condition (Fig. 3A, B). Sequencing verified the presence of restored full-length eGFP RNA containing the expected junctions at Csm cleavage positions (Fig 3B, Supplementary Table 6).

We hypothesized that increasing the expression and stability of our reporter transcripts would result in more opportunities for *trans*-ligation and more reconstituted eGFP protein. We therefore increased transcriptional output by switching to the CAG promoter, and also increased the half-life of the donor and subsequent *trans*-ligated product by installing a Woodchuck Hepatitis Virus Post-transcriptional Regulatory Element (WPRE) and the bovine growth hormone polyadenylation signal (BGHpA) at the 3′ end of the transcript encoding the C-terminal eGFP fragment (Fig 3C). Strikingly, this optimized reporter system (Donor-v2) dramatically increased the fraction of eGFP+ cells to over 40% as well as their fluorescence intensity only when a targeting crRNA was expressed, indicating successful *trans*-ligation (Fig. 3C, Fig. S2). We term the *in vivo* production of chimeric RNAs mediated by Csm “spligation” to distinguish it from spliceosome-dependent *trans*-splicing techniques.

Finally, we explored alternate methods to generate donor transcripts possessing the required 5′-hydroxyl terminus. In place of the Csm target site on the donor transcript, we encoded one of several *cis*-cleaving ribozymes, or a Csm crRNA stem loop, which is recognized and cleaved by the Cas6 already encoded in our Csm plasmid (Fig 3D). Indeed, eGFP fluorescence intensity in all conditions where the donor 5′-hydroxyl was generated by ribozymes or Cas6 was 2-3 fold higher than with the standard 2-cut system (Fig 3D). As expected, these alternate donor transcripts (Donor-v3) also led to reduced mCherry fluorescence intensity even in the absence of a targeting crRNA, supporting the efficient separation of the 3′ end. Additionally, these alternative cleavage mechanisms enable donor transcripts with precise 5′ identities, in principle reducing variation in the edited products from multiple Csm cut sites. These results offer a unique opportunity to pair site-specific transcript cleavage with chemically compatible donor RNA molecules.

### Spligation enables splice-site independent endogenous mRNA modification

Modification of endogenous transcripts in such a precise manner has been limited to classical *trans*-splicing, which relies on natural intron-exon boundaries. Given the success of spligation in our reporter assay, we sought to determine whether Csm could mediate site-specific spligation to endogenous mRNA molecules. Of the alternate donor cleavage mechanisms we tested, all but the RzB hammerhead ribozyme require inclusion of extra nucleotides required for secondary structure at the 5′ end of the cleaved transcript. Therefore, we redesigned our Donor-v3 plasmid to encode the Rzb hammerhead ribozyme just upstream of a glycine-serine rich linker followed by full-length superfolder GFP (sfGFP) as a C-terminal protein tag (Fig. 4A).

**Figure 4:**
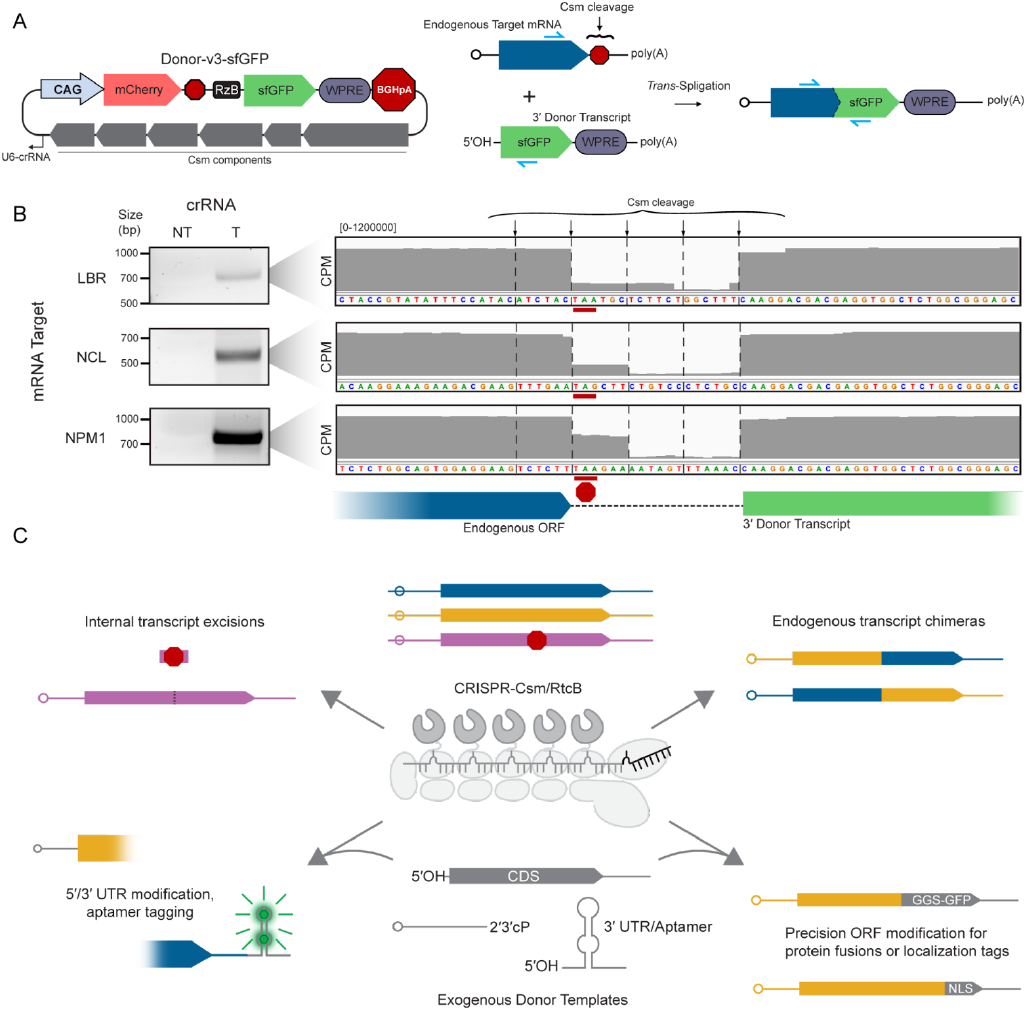
Spligation with Csm enables spliceosome-independent multi-kilobase additions to endogenous mRNA transcripts. **(A)** Schematic of Donor-v3 plasmid equipped with RzB hammerhead ribozyme repurposed for spligation to endogenous transcripts (left). Csm cleavage targeted to the stop codon of endogenous protein-coding genes results in the addition of sfGFP to the open reading frame (right). **(B)** RNA was extracted from cells transfected with the constructs outlined in (A), reverse-transcribed into cDNA, and PCR amplified using the primers indicated in blue in (A). The PCR product was resolved on a 1.5% agarose gel and the indicated band excised and sequenced. Insets (right) show sequencing coverage across the Csm cleavage sites for each target. Reporter sequences and diagrams depicted below. **(C)** Visual summary of spligation as a versatile method for transcriptome engineering *in vivo*, highlighting current and future applications.

We targeted Csm to one of three unique endogenous mRNAs, aligning the crRNA to locate the predominant scissile phosphate at the 5′ end of the stop codon. Following transfection of HEK293T cells with the Donor-v3 plasmid, we were able to verify chimeric transcripts by RT-PCR for all targets tested, confirming spligation of the donor molecule to the endogenous transcript (Fig. 4B, Supplementary Tables 7-9). These results constitute, to our knowledge, the first report of fully site-programmable “*trans*-splicing” to endogenous transcripts relying on a cut-and-paste mechanism rather than natural or engineered intron-exon boundaries.

## Discussion

Based on observations made during the use of CRISPR-Csm for targeted ablation of cellular transcripts, we identified two forms of precision RNA manipulation that can be induced in live human cells. By introducing CRISPR-Csm complexes and suitable guide RNAs into cells, site-specific transcript cleavage can be repaired by RNA ligation pathways, introducing excisions of at least several hundred nucleotides within an mRNA transcript without disrupting the reading frame. Csm-mediated cleavage of two different transcripts can generate *trans*-ligated chimeric RNAs, or spligation products, not delimited by naturally occurring splice sites. Together with ligation-enhancing CRISPR tools, these new forms of RNA manipulation may unlock previously inaccessible methods of exploring RNA biology.

In eukaryotes, endonucleolytic RNA cleavage usually results in transcript-wide degradation by host 5′-3′ and 3′-5′ exonucleases^16^. Thus, it was striking to observe a small but significant fraction of Csm-cleaved transcripts that had been re-ligated, as evidenced by the distinctive 6-nt incremental footprint of missing sequences. While there are two known examples of RNA cleavage and re-ligation in eukaryotes outside of the spliceosome (tRNA maturation and unconventional splicing of XBP1)^10^, both catalyzed by the RNA ligase RtcB, this enzyme was not known to act on other endogenous RNA substrates. It will be interesting to see if RtcB ligates other transcripts or naturally occurring cleavage products in the cell, and with what degree of specificity.

The reporter assay developed in this study allows detection of both Csm-mediated cleavage and RtcB-mediated repair by monitoring reduction of mCherry fluorescence and appearance of eGFP fluorescence, respectively. The low efficiency of RNA ligation observed in cells could be enhanced by direct fusion of RtcB and Csm3. Interestingly, bacterial RtcB performed better than the human ortholog, perhaps due to its cofactor-free, multiple-turnover activity^17^. Additional engineering efforts or biological insights into RNA repair pathways will be critical to further favor religation efficiency over transcript degradation.

A natural advantage of the Csm complex over Cas13 nucleases is its ability to cut RNA site-specifically and only in *cis*^18^. Additionally, Csm is unique in its characteristic cleavage pattern that always occurs in multiples of 6 nts across the target site. This characteristic activity allows short segments of RNA to be excised with a single cleavage event in a manner that always maintains reading frame, a property we and others^7,19^ exploited to remove nonsense mutations. Importantly, following RNA excision, the target recognition sequence is destroyed, rendering edited transcripts immune to further cleavage by Csm. Thus, this method allows for clearance of unwanted transcripts and enrichment of edited ones, demonstrated by the robust gene expression in our reporter system.

Beyond short excisions (microexcisions) via single cleavage events, we and others^7^ demonstrated Csm’s utility for generating longer deletions (macroexcisions) by cleaving multiple times within the same transcript. Specifically, we removed up to ∼300 nts of sequence, substantially longer than what is achievable by single-cleavage events (6-24 nts). This approach could in principle be extended to remove entire exons (e.g. poison exons) or toxic repeat expansions for therapeutic purposes, by cutting on either side of the unwanted element with multiplexed guide RNAs.

Finally, by cleaving two separate transcripts, spligation produced unique RNAs stitched together into a single chimeric fusion. Altered mRNA sequences can be tested from artificial expression from a plasmid or viral vector, but these methods do not necessarily recapitulate the endogenous genomic context and natural epigenetic regulation. Modifications to endogenous transcripts typically require either providing a complete new exon(s) or genome editing to make the desired change. Spligation can be used to generate new RNA isoforms or replace regions of a transcript with exogenously delivered RNA templates at virtually any position, without requiring time-consuming genome editing or artificial overexpression. Spligation could therefore have a variety of applications in fundamental research (Fig. 4D). For example, tagging or replacing 5′ or 3′ transcript ends could alter protein products or untranslated regions without affecting natural transcriptional regulation. Chimeric transcripts could also enable direct monitoring of transcripts using fluorogenic aptamers, reversion of transcription products following chromosomal translocations, or genome editing of RNA viruses within host cells. Spligation expands the RNA editing toolkit, providing a new, fully programmable option for manipulating eukaryotic transcripts free of exonic restraints.

## Materials and Methods

### Cell culture

HEK293T cells (UC Berkeley Cell Culture Facility) were grown in medium containing DMEM, high glucose, GlutaMAX supplement, sodium pyruvate (Thermo Fisher Scientific), 10% FBS (Sigma), 25 mM HEPES pH 7.2–7.5 (Thermo Fisher Scientific), 1× MEM nonessential amino acids (Thermo Fisher Scientific), 1× Pen/Strep (Thermo Fisher Scientific) and 0.1 mM βME (Thermo Fisher Scientific) at 37 °C with 5% CO2. All cell lines were verified to be mycoplasma-free (abm, PCR mycoplasma detection kit).

### Plasmid Construction

CRISPR-Csm plasmid was previously described11 (pDAC435, Addgene #195238). Reporter constructs (cis and trans-ligation reporters and endogenous transcript donors) were installed between the CMV::Cas6 and the U6::crRNA expression cassettes. RtcB sequences from *Escherichia coli, Homo sapiens*, and *Pyrococcus horikoshii* were obtained from NCBI. Protein sequences were human codon optimized using online tools (GenScript), synthesized as gene blocks (IDT), and cloned as Csm3-RtcB fusions using ClaI and SpeI restriction sites. Plasmids were verified by whole-plasmid sequencing. Plasmid sequences are provided in Supplementary Table 10.

### DNA transfections

Following the manufacturer’s instructions, 1 × 106 HEK293T cells were transfected with 2 µg plasmid DNA using 6 µl FuGENE HD transfection reagent in 6-well plates. Following transfection, cells were grown for 48 h to allow plasmid expression prior to assaying.

### Flow cytometry

Cell fluorescence was assayed on an Attune NxT acoustic focusing cytometer (Thermo Fisher Scientific) equipped with 488 nm excitation laser and 530/30 emission filter (eGFP), and 561 nm excitation laser and 620/15 emission filter (mCherry). Data were analyzed using Attune Cytometric Software v5.1.1 and FlowJo v10.7.1.

### RT-PCR

Total cellular RNA was extracted using TRIzol Reagent (Thermo Fisher Scientific) as per the manufacturer’s instructions. Genomic DNA was removed using TURBO DNase (Thermo Fisher Scientific). After inactivating TURBO DNase with DNase Inactivating Reagent, 1 µg DNase-free RNA was reverse transcribed into cDNA using Induro Reverse Transcriptase (NEB) with oligo(dT) primers (Promega) as per the manufacturer’s instructions. PCR was performed on cDNA template using Q5 High-Fidelity 2x Master Mix (NEB) in a ProFlex PCR System (Applied Biosystems). Gene-specific primer pairs are listed in Supplementary Table 11. No-RT and no-template controls were run alongside all RT-PCR experiments.

### Nanopore sequencing

PCR amplicons from cDNA were separated on a 1.5% agarose gel, excised and purified using QIAquick Gel Extraction Kit (Qiagen). Samples were sequenced using Oxford Nanopore Technology (Plasmidsaurus). Each product’s composition was confirmed by aligning reads to the corresponding amplicon with minimap2 (v2.28) and a custom junction file representing Csm cut sites. The abundance of each excision or ligation product was determined with HTSlib (v1.6)^20^, samtools (v1.6)^21^, and pysam (v0.18.0)^22^ and reported relative to total read depth within the crRNA target site.

### RNA-seq analysis

RNA-data were re-analyzed from our previous work (Colognori et al, 2023; GSE220741)^11^. Ligation efficiency was quantified using a custom Python (version 3.9.12) script. Briefly, FASTQ files were aligned to the GRCh38 reference genome (GENCODE Release 3923) with STAR (v2.7.10a)^24^. The number of reads corresponding to individual XIST ligation products was quantified using HTSlib (v1.6)^20^, samtools (v1.6)^21^, and pysam (v0.18.0)^22^, and normalized to the total number of reads spanning the target site.

### Microscopy

For wide-field fluorescence imaging, live cells were observed in 6-well plates on an EVOS FL inverted fluorescence microscope (Thermo Fisher), equipped with LPlanFL PH2 10x/0.30NA objective lens, GFP 2.0 (482/25 ex, 524/24 em) and Texas Red 2.0 (585/29 ex, 628/32 em) light cubes, and Sony ICX445 monochrome CCD camera (1280 x 960 pixel resolution, 1.3 Megapixels). Image analysis was performed using FIJI (ImageJ) software.

### Statistical analysis

All graphs display the mean and standard deviation of three biological replicates. For RNA-seq analysis, no statistical parameters were applied, given one biological replicate.

## Data and materials availability

All data will be released upon publication. Plasmids corresponding to this work will be made available on Addgene and raw sequencing data will be deposited on GEO and SRA.

## References

1. Jinek, M. et al. A programmable dual-RNA-guided DNA endonuclease in adaptive bacterial immunity. Science 337, 816–821 (2012).

2. Mali, P. et al. RNA-guided human genome engineering via Cas9. Science 339, 823–826 (2013).

3. Abudayyeh, O. O. et al. C2c2 is a single-component programmable RNA-guided RNA-targeting CRISPR effector. Science 353, aaf5573 (2016).

4. Ai, Y., Liang, D. & Wilusz, J. E. CRISPR/Cas13 effectors have differing extents of off-target effects that limit their utility in eukaryotic cells. Nucleic Acids Res. 50, e65 (2022).

5. Shi, P. et al. Collateral activity of the CRISPR/RfxCas13d system in human cells. Commun. Biol. 6, 334 (2023).

6. Fiflis, D. N. et al. Repurposing CRISPR-Cas13 systems for robust mRNA trans-splicing. Nat. Commun. 15, 2325 (2024).

7. Nemudraia, A., Nemudryi, A. & Wiedenheft, B. Repair of CRISPR-guided RNA breaks enables site-specific RNA excision in human cells. Science 384, 808–814 (2024).

8. Irmisch, P., Mogila, I., Samatanga, B., Tamulaitis, G. & Seidel, R. Retention of the RNA ends provides the molecular memory for maintaining the activation of the Csm complex. Nucleic Acids Res. 52, 3896–3910 (2024).

9. Moncan, M., Rakhsh-Khorshid, H., Eriksson, L. A., Samali, A. & Gorman, A. M. Insights into the structure and function of the RNA ligase RtcB. Cell Mol Life Sci 80, 352 (2023).

10. Lu, Y., Liang, F.-X. & Wang, X. A synthetic biology approach identifies the mammalian UPR RNA ligase RtcB. Mol. Cell 55, 758–770 (2014).

11. Colognori, D., Trinidad, M. & Doudna, J. A. Precise transcript targeting by CRISPR-Csm complexes. Nat. Biotechnol. 41, 1256–1264 (2023).

12. Tamulaitis, G. et al. Programmable RNA shredding by the type III-A CRISPR-Cas system of Streptococcus thermophilus. Mol. Cell 56, 506–517 (2014).

13. Xia, C., Colognori, D., Jiang, X. S., Xu, K. & Doudna, J. A. Single-molecule live-cell RNA imaging with CRISPR–Csm. Nature Biotechnology 43, 2023–2030 (2025).

14. Lindley, S. R. et al. Ribozyme-activated mRNA trans-ligation enables large gene delivery to treat muscular dystrophies. Science 386, 762–767 (2024).

15. Ghosh, I., Hamilton, A. D. & Regan, L. Antiparallel leucine zipper-directed protein reassembly: Application to the Green fluorescent protein. J. Am. Chem. Soc. 122, 5658–5659 (2000).

16. Houseley, J. & Tollervey, D. The many pathways of RNA degradation. Cell 136, 763–776 (2009).

17. Desai, K. K., Beltrame, A. L. & Raines, R. T. Coevolution of RtcB and Archease created a multiple-turnover RNA ligase. RNA 21, 1866–1872 (2015).

18. Staals, R. H. J. et al. RNA targeting by the type III-A CRISPR-Cas Csm complex of Thermus thermophilus. Mol. Cell 56, 518–530 (2014).

19. Sun, Y. et al. Type III CRISPR-mediated flexible RNA excision with engineered guide RNAs. Mol. Cell 85, 989–998.e4 (2025).

20. Bonfield, J. K. et al. HTSlib: C library for reading/writing high-throughput sequencing data. Gigascience 10, (2021).

21. Danecek, P. et al. Twelve years of SAMtools and BCFtools. Gigascience 10, (2021).

22. Heger, A. et al. pysam (v0.18.0). GitHub (2023). https://github.com/pysam-developers/pysam

23. Frankish, A. et al. GENCODE: reference annotation for the human and mouse genomes in 2023. Nucleic Acids Res. 51, D942–D949 (2023).

24. Dobin, A. et al. STAR: ultrafast universal RNA-seq aligner. Bioinformatics 29, 15–21 (2013).

