## Supplementary_Figures_and_Tables for "Spligation enables programmable chimeric RNA generation in living cells"

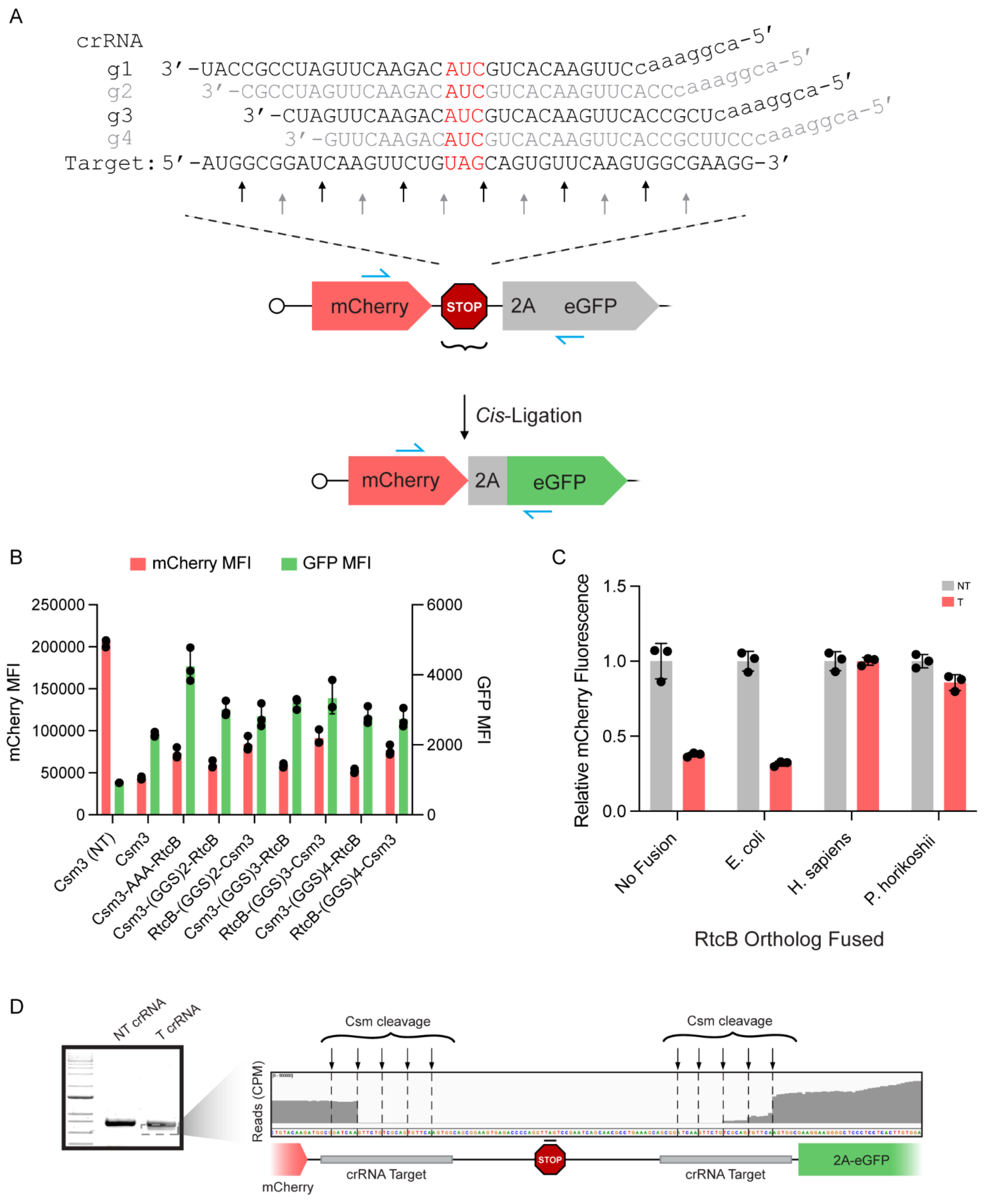

**Supplementary Fig 1**

**A.** Positioning of tiled crRNA sequences relative to the reporter construct stop codon region, corresponding to Figure 2C. Black arrows indicate possible cleavage sites for crRNAs 1 & 3, while gray arrows indicate possible cleavage sites for crRNAs 2 & 4.

**B.** Flow cytometry results from experiment testing fusion orientations & linker lengths.
C-terminal Csm3 fusion with AAA linker performed the best and was used in all subsequent experiments.

**C.** Flow cytometry quantification of mCherry knockdown in RtcB fusion comparison experiment, corresponding to Figure 2E.

**D.** RNA was extracted from cells transfected with the construct shown in Fig. **2G** with the crRNA target sites spaced ~50 nt apart, reverse-transcribed into cDNA, and PCR amplified using the primers indicated in blue in (**Fig. 2G**). The PCR product was resolved on a 1.5% agarose gel and the indicated band excised and sequenced. Inset shows sequencing coverage across the Csm cleavage sites. Reporter sequence and diagram depicted below.

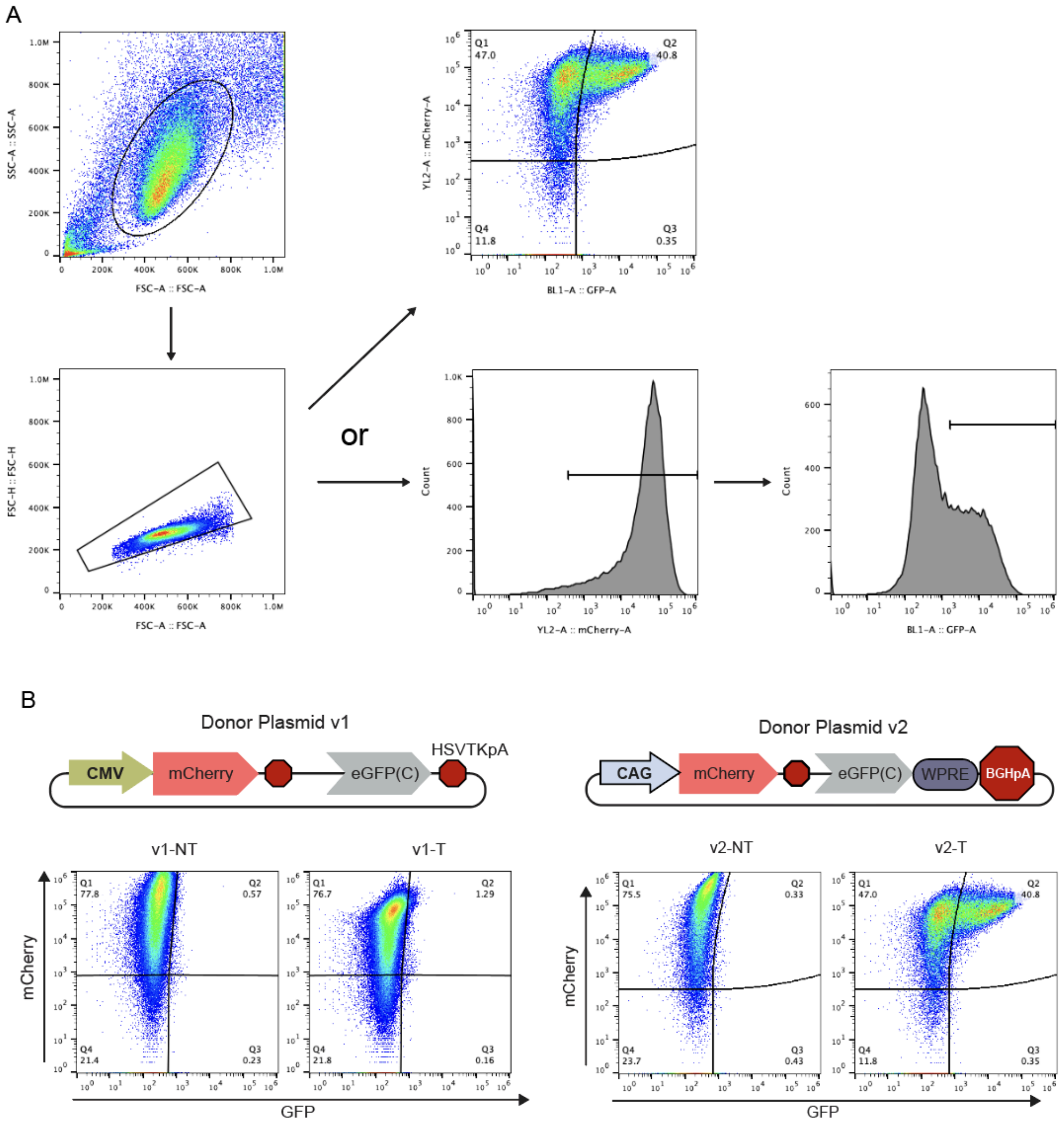


**Supplementary Fig 2**

**A.** Gating strategy for flow cytometry analysis, representative of all experiments.

**B.** Representative flow cytometry panels for trans-ligation experiments using original Donor-v1 (top) and enhanced Donor-v2 (bottom) constructs. Cells transfected with a non-targeting crRNA shown on left, targeting crRNA on right.

| Ligation Product: XIST crRNA 2 | N_Reads | Percent |
|---|---|---|
| CATGGCGGGCTGTGCTTTGTTAGGTTGTCCAA | 550 | 75.86206897 |
| CATGGCGGGCTGTGC------AGGTTGTCCAA | 32 | 4.413793103 |
| CATGGCGGG------TTTGTTAGGTTGTCCAA | 30 | 4.137931034 |
| CATGGCGGG------------AGGTTGTCCAA | 21 | 2.896551724 |
| CAT------------TTTGTTAGGTTGTCCAA | 4 | 0.551724138 |
| CATGGCGGG------------------TCCAA | 2 | 0.275862069 |
| CAT------------------AGGTTGTCCAA | 1 | 0.137931034 |
| CAT------CTGTGCTTTGTTAGGTTGTCCAA | 0 | 0 |
| CAT------------------------TCCAA | 0 | 0 |
| CATGGCGGGCTGTGC------------TCCAA | 0 | 0 |
| CATGGCGGGCTGTGCTTTGTT------TCCAA | 0 | 0 |
| Other_Reads | 85 | 11.72413793 |
| Total_Reads | 725 | 100 |

**Supplementary Table 1**

| Ligation Product: XIST crRNA 1 | N_Reads | Percent |
|---|---|---|
| CGGATCCAGTTCTGTCGCAGTGTTCAAGTGGC | 283 | 76.90217391 |
| CGGATCCAG------------GTTCAAGTGGC | 15 | 4.076086957 |
| CGGATCCAGTTCTGT------GTTCAAGTGGC | 9 | 2.445652174 |
| CGGATCCAGTTCTGT------------GTGGC | 14 | 3.804347826 |
| CGGATCCAG------CGCAGTGTTCAAGTGGC | 9 | 2.445652174 |
| CGGATCCAG------------------GTGGC | 7 | 1.902173913 |
| CGG------TTCTGTCGCAGTGTTCAAGTGGC | 0 | 0 |
| CGG------------CGCAGTGTTCAAGTGGC | 0 | 0 |
| CGG------------------GTTCAAGTGGC | 0 | 0 |
| CGG------------------------GTGGC | 0 | 0 |
| CGGATCCAGTTCTGTCGCAGT------GTGGC | 0 | 0 |
| Other_Reads | 31 | 8.423913043 |
| Total_Reads | 368 | 100 |

**Supplementary Table 2**

| **Ligation Product: Cis 1-Cut Reporter** | **N_Reads** | **Percent** |
|---|---|---|
| GCGGATCAA------------------AGTGG | 475 | 64.4504749 |
| GCGGATCAAGTTCTGTAGCAGTGTTCAAGTGG | 30 | 4.070556309 |
| GCGGATCAA------TAGCAGTGTTCAAGTGG | 56 | 7.598371777 |
| GCGGATCAA------------TGTTCAAGTGG | 24 | 3.256445047 |
| GCGGATCAAGTTCTG------------AGTGG | 15 | 2.035278155 |
| GCG------------------------AGTGG | 6 | 0.814111262 |
| GCG------GTTCTGTAGCAGTGTTCAAGTGG | 0 | 0 |
| GCG------------TAGCAGTGTTCAAGTGG | 0 | 0 |
| GCG------------------TGTTCAAGTGG | 0 | 0 |
| GCGGATCAAGTTCTG------TGTTCAAGTGG | 0 | 0 |
| GCGGATCAAGTTCTGTAGCAG------AGTGG | 0 | 0 |
| Other_Reads | 131 | 17.77476255 |
| Total_Reads | 737 | 100 |

**Supplementary Table 3**

| **Ligation Product: 2-Cut 100nt Reporter** | **N_Reads** | **Percent** |
|---|---|---|
| GCGGATCAA-----------------------\|----------------<br>-----------AGTGG | 205 | 65.2866242 |
| GCG-----------------------------\|----------------<br>-----------AGTGG | 1 | 0.318471338 |
| GCGGATCAA-----------------------\|----------------<br>-----TGTTCAAGTGG | 11 | 3.503184713 |
| GCGGATCAA-----------------------\|---------------T<br>CGCAGTGTTCAAGTGG | 21 | 6.687898089 |
| GCGGATCAAGTTCTGTCGCAG-----------\|----------------<br>-----------AGTGG | 1 | 0.318471338 |
| GCGGATCAAGTTCTGTCGCAG-----------\|----------------<br>-----TGTTCAAGTGG | 5 | 1.592356688 |
| GCG-----------------------------\|---GATCAAGTTCTGT<br>CGCAGTGTTCAAGTGG | 0 | 0 |
| GCG-----------------------------\|---------GTTCTGT<br>CGCAGTGTTCAAGTGG | 0 | 0 |
| GCG-----------------------------\|---------------T<br>CGCAGTGTTCAAGTGG | 0 | 0 |
| GCG-----------------------------\|----------------<br>-----TGTTCAAGTGG | 0 | 0 |
| GCGGATCAA-----------------------\|---GATCAAGTTCTGT<br>CGCAGTGTTCAAGTGG | 0 | 0 |
| GCGGATCAA-----------------------\|---------GTTCTGT<br>CGCAGTGTTCAAGTGG | 0 | 0 |
| GCGGATCAAGTTCTG-----------------\|---GATCAAGTTCTGT<br>CGCAGTGTTCAAGTGG | 0 | 0 |
| GCGGATCAAGTTCTG-----------------\|---------GTTCTGT<br>CGCAGTGTTCAAGTGG | 0 | 0 |
| GCGGATCAAGTTCTG-----------------\|---------------T<br>CGCAGTGTTCAAGTGG | 0 | 0 |
| GCGGATCAAGTTCTG-----------------\|---------------- | 0 | 0 |

| | | |
|---|---|---|
| -----TGTTCAAGTGG |  |  |
| GCGGATCAAGTTCTG-----------------\|---------------- -----------AGTGG | 0 | 0 |
| GCGGATCAAGTTCTGTCGCAG-----------\|---GATCAAGTTCTGT CGCAGTGTTCAAGTGG | 0 | 0 |
| GCGGATCAAGTTCTGTCGCAG-----------\|---------GTTCTGT CGCAGTGTTCAAGTGG | 0 | 0 |
| GCGGATCAAGTTCTGTCGCAG-----------\|---------------T CGCAGTGTTCAAGTGG | 0 | 0 |
| GCGGATCAAGTTCTGTCGCAGTGTTCA-----\|---GATCAAGTTCTGT CGCAGTGTTCAAGTGG | 0 | 0 |
| GCGGATCAAGTTCTGTCGCAGTGTTCA-----\|---------GTTCTGT CGCAGTGTTCAAGTGG | 0 | 0 |
| GCGGATCAAGTTCTGTCGCAGTGTTCA-----\|---------------T CGCAGTGTTCAAGTGG | 0 | 0 |
| GCGGATCAAGTTCTGTCGCAGTGTTCA-----\|---------------- -----TGTTCAAGTGG | 0 | 0 |
| GCGGATCAAGTTCTGTCGCAGTGTTCA-----\|---------------- -----------AGTGG | 0 | 0 |
| Other_Reads | 70 | 22.29299363 |
| Total_Reads | 314 | 100 |

**Supplementary Table 4**

| Ligation Product: 2-Cut 300nt Reporter | N_Reads | Percent |
|---|---|---|
| GCGGATCAA-----------------------\|---------------- -----------AGTGG | 148 | 48.05194805 |
| GCGGATCAA-----------------------\|---------------T CGCAGTGTTCAAGTGG | 48 | 15.58441558 |
| GCGGATCAA-----------------------\|---------------- -----TGTTCAAGTGG | 6 | 1.948051948 |
| GCGGATCAAGTTCTGTCGCAG-----------\|---------------- -----TGTTCAAGTGG | 34 | 11.03896104 |
| GCG-----------------------------\|---------------- -----------AGTGG | 1 | 0.324675325 |
| GCG-----------------------------\|---GATCAAGTTCTGT CGCAGTGTTCAAGTGG | 0 | 0 |
| GCG-----------------------------\|---------GTTCTGT CGCAGTGTTCAAGTGG | 0 | 0 |
| GCG-----------------------------\|---------------T CGCAGTGTTCAAGTGG | 0 | 0 |
| GCG-----------------------------\|---------------- -----TGTTCAAGTGG | 0 | 0 |
| GCGGATCAA-----------------------\|---GATCAAGTTCTGT CGCAGTGTTCAAGTGG | 0 | 0 |
| GCGGATCAA-----------------------\|---------GTTCTGT CGCAGTGTTCAAGTGG | 0 | 0 |
| GCGGATCAAGTTCTG-----------------\|---GATCAAGTTCTGT CGCAGTGTTCAAGTGG | 0 | 0 |
| GCGGATCAAGTTCTG-----------------\|---------GTTCTGT CGCAGTGTTCAAGTGG | 0 | 0 |
| GCGGATCAAGTTCTG-----------------\|---------------T CGCAGTGTTCAAGTGG | 0 | 0 |
| GCGGATCAAGTTCTG-----------------\|---------------- -----TGTTCAAGTGG | 0 | 0 |
| GCGGATCAAGTTCTG-----------------\|---------------- | 0 | 0 |

| | | |
|---|---|---|
| -----------AGTGG |  |  |
| GCGGATCAAGTTCTGTCGCAG-----------\|---GATCAAGTTCTGT<br>CGCAGTGTTCAAGTGG | 0 | 0 |
| GCGGATCAAGTTCTGTCGCAG-----------\|---------GTTCTGT<br>CGCAGTGTTCAAGTGG | 0 | 0 |
| GCGGATCAAGTTCTGTCGCAG-----------\|---------------T<br>CGCAGTGTTCAAGTGG | 0 | 0 |
| GCGGATCAAGTTCTGTCGCAG-----------\|----------------<br>-----------AGTGG | 0 | 0 |
| GCGGATCAAGTTCTGTCGCAGTGTTCA-----\|---GATCAAGTTCTGT<br>CGCAGTGTTCAAGTGG | 0 | 0 |
| GCGGATCAAGTTCTGTCGCAGTGTTCA-----\|---------GTTCTGT<br>CGCAGTGTTCAAGTGG | 0 | 0 |
| GCGGATCAAGTTCTGTCGCAGTGTTCA-----\|---------------T<br>CGCAGTGTTCAAGTGG | 0 | 0 |
| GCGGATCAAGTTCTGTCGCAGTGTTCA-----\|----------------<br>-----TGTTCAAGTGG | 0 | 0 |
| GCGGATCAAGTTCTGTCGCAGTGTTCA-----\|----------------<br>-----------AGTGG | 0 | 0 |
| Other_Reads | 71 | 23.05194805 |
| Total_Reads | 308 | 100 |

**Supplementary Table 5**

| Ligation Product: Trans-Ligation Reporter | N_Reads | Percent |
|---|---|---|
| GCGGATCAA-----------------------\|---------------TCGCAGTGTTCAAGTGG | 1949 | 40.72294191 |
| GCGGATCAA-----------------------\|---------------------TGTTCAAGTGG | 364 | 7.605516089 |
| GCGGATCAA-----------------------\|---------------------------AGTGG | 1863 | 38.92603427 |
| GCG-----------------------------\|---------------------------AGTGG | 13 | 0.271625575 |
| GCGGATCAAGTTCTG-----------------\|---------------------TGTTCAAGTGG | 3 | 0.062682825 |
| GCGGATCAAGTTCTG-----------------\|---------GTTCTGTCGCAGTGTTCAAGTGG | 3 | 0.062682825 |
| GCG-----------------------------\|---------------------TGTTCAAGTGG | 1 | 0.020894275 |
| GCG-----------------------------\|---GATCAAGTTCTGTCGCAGTGTTCAAGTGG | 0 | 0 |
| GCG-----------------------------\|---------GTTCTGTCGCAGTGTTCAAGTGG | 0 | 0 |
| GCG-----------------------------\|---------------TCGCAGTGTTCAAGTGG | 0 | 0 |
| GCGGATCAA-----------------------\|---GATCAAGTTCTGTCGCAGTGTTCAAGTGG | 0 | 0 |
| GCGGATCAA-----------------------\|---------GTTCTGTCGCAGTGTTCAAGTGG | 0 | 0 |
| GCGGATCAAGTTCTG-----------------\|---GATCAAGTTCTGTCGCAGTGTTCAAGTGG | 0 | 0 |
| GCGGATCAAGTTCTG-----------------\|---------------TCGCAGTGTTCAAGTGG | 0 | 0 |
| GCGGATCAAGTTCTG-----------------\|---------------------------AGTGG | 0 | 0 |
| GCGGATCAAGTTCTGTCGCAG-----------\|---GATCAAGTTCTG | 0 | 0 |

| | | |
|---|---|---|
| TCGCAGTGTTCAAGTGG | | |
| GCGGATCAAGTTCTGTCGCAG-----------\|---------GTTCTG TCGCAGTGTTCAAGTGG | 0 | 0 |
| GCGGATCAAGTTCTGTCGCAG-----------\|--------------- TCGCAGTGTTCAAGTGG | 0 | 0 |
| GCGGATCAAGTTCTGTCGCAG-----------\|--------------- ------TGTTCAAGTGG | 0 | 0 |
| GCGGATCAAGTTCTGTCGCAG-----------\|--------------- ------------AGTGG | 0 | 0 |
| GCGGATCAAGTTCTGTCGCAGTGTTCA-----\|---GATCAAGTTCTG TCGCAGTGTTCAAGTGG | 0 | 0 |
| GCGGATCAAGTTCTGTCGCAGTGTTCA-----\|---------GTTCTG TCGCAGTGTTCAAGTGG | 0 | 0 |
| GCGGATCAAGTTCTGTCGCAGTGTTCA-----\|--------------- TCGCAGTGTTCAAGTGG | 0 | 0 |
| GCGGATCAAGTTCTGTCGCAGTGTTCA-----\|--------------- ------TGTTCAAGTGG | 0 | 0 |
| GCGGATCAAGTTCTGTCGCAGTGTTCA-----\|--------------- ------------AGTGG | 0 | 0 |
| Other_Reads | 590 | 12.32762223 |
| Total_Reads | 4786 | 100 |

**Supplementary Table 6**

| Ligation Product: LBR | N_Reads | Percent |
|---|---|---|
| TACATCTAC------------------ | 3153 | 81.2628866 |
| TACATCTACTAATGCTCTTCT------ | 549 | 14.14948454 |
| TAC------TAATGCTCTTCTGGCTTT | 0 | 0 |
| TAC------------TCTTCTGGCTTT | 0 | 0 |
| TAC------------------GGCTTT | 0 | 0 |
| TAC------------------------ | 0 | 0 |
| TACATCTAC------TCTTCTGGCTTT | 0 | 0 |
| TACATCTAC------------GGCTTT | 0 | 0 |
| TACATCTACTAATGC------GGCTTT | 0 | 0 |
| TACATCTACTAATGC------------ | 0 | 0 |
| Other_Reads | 178 | 4.587628866 |
| Total_Reads | 3880 | 100 |

**Supplementary Table 7**

| Ligation Product: NCL | N_Reads | Percent |
|---|---|---|
| AAGTTTGAA------------------ | 2956 | 61.686 |
| AAGTTTGAATAGCTT------------ | 967 | 20.179 |
| AAG------------------------ | 73 | 1.5234 |
| AAG------TAGCTTCTGTCCCTCTGC | 0 | 0 |
| AAG------------CTGTCCCTCTGC | 0 | 0 |
| AAG------------------CTCTGC | 0 | 0 |
| AAGTTTGAA------CTGTCCCTCTGC | 0 | 0 |
| AAGTTTGAA------------CTCTGC | 0 | 0 |
| AAGTTTGAATAGCTT------CTCTGC | 0 | 0 |
| AAGTTTGAATAGCTTCTGTCC------ | 0 | 0 |
| Other_Reads | 796 | 16.611 |
| Total_Reads | 4792 | 100 |

**Supplementary Table 8**

| Ligation Product: NPM1 | N_Reads | Percent |
|---|---|---|
| AAGTCTCTTTAAGAA------------ | 2082 | 43.17710494 |
| AAGTCTCTT------------------ | 1827 | 37.8888428 |
| AAG------TAAGAAAATAGTTTAAAC | 0 | 0 |
| AAG------------AATAGTTTAAAC | 0 | 0 |
| AAG------------------TTAAAC | 0 | 0 |
| AAG------------------------ | 0 | 0 |
| AAGTCTCTT------AATAGTTTAAAC | 0 | 0 |
| AAGTCTCTT------------TTAAAC | 0 | 0 |
| AAGTCTCTTTAAGAA------TTAAAC | 0 | 0 |
| AAGTCTCTTTAAGAAAATAGT------ | 0 | 0 |
| Other_Reads | 913 | 18.93405226 |
| Total_Reads | 4822 | 100 |

**Supplementary Table 9**
